# Comparison of AAV9-driven motor neuron transduction following different CNS-directed delivery methods in mice

**DOI:** 10.1101/2025.07.01.662642

**Authors:** Alannah J Mole, Chiara F Sander, Amisha R Parmar, Ailsa J Williams, Mimoun Azzouz, Guillaume M Hautbergue, Pamela J Shaw, Laura Ferraiuolo, Richard J Mead

## Abstract

**Background:** Gene therapies are promising for diseases previously considered incurable. Adeno-associated virus serotype 9 (AAV9) demonstrates remarkable tropism for motor neurons (MNs) and represents an exciting candidate to target genetic causes of motor neuron diseases like amyotrophic lateral sclerosis (ALS). However, systemic delivery risks immunogenicity and off-target effects, therefore localised delivery to the CNS is advantageous.

**New method:** We assessed MN transduction in wild-type mice using AAV9-controlled, cytomegalovirus-promoter driven, enhanced GFP expression. Intra-cisterna magna (ICM) and intra-cerebroventricular (ICV) methods were compared. Four weeks post-delivery, GFP positivity in MN and astrocytes were quantified via immunohistochemical approaches and viral genome copy number determined by qPCR.

**Results:** All delivery methods achieved high MN transduction in lumbar spinal cord (>68%). Unilateral ICV delivery provided the highest and most consistent levels (89 ± 3%), and minimal peripheral viral copies. ICV delivery resulted in higher astrocytic transduction, most notably in the cortex. Brainstem MN transduction was high with all methods (>55%). We failed to find evidence of neuronal transduction in motor cortex. Viral genome copies trended higher in spinal cord and brainstem with ICV approaches, however further work is required to understand how bilateral delivery leads to such profound increases.

**Comparison to existing methods:** Whilst several routes of administration into cerebrospinal fluid exist, direct comparisons for targeting MNs *in vivo* remain limited.

**Conclusions:** Overall, consideration of gene therapy delivery methods is critical to ensure that the most appropriate administration route is chosen to reach MNs effectively, selectively, and at high levels to exact biological effects.

**Highlights:**

- CSF delivery of AAV9 targets >68% of lumbar spinal cord (SC) motor neurons
- Unilateral ICV yields high and consistent lumbar SC motor neuron transduction (>88%)
- Unilateral ICV results in the lowest number of peripheral viral copies
- ICV targets significantly more astrocytes, particularly in cortex
- Bilateral ICV leads to significantly higher viral copies in the liver

**Graphical Abstract:** 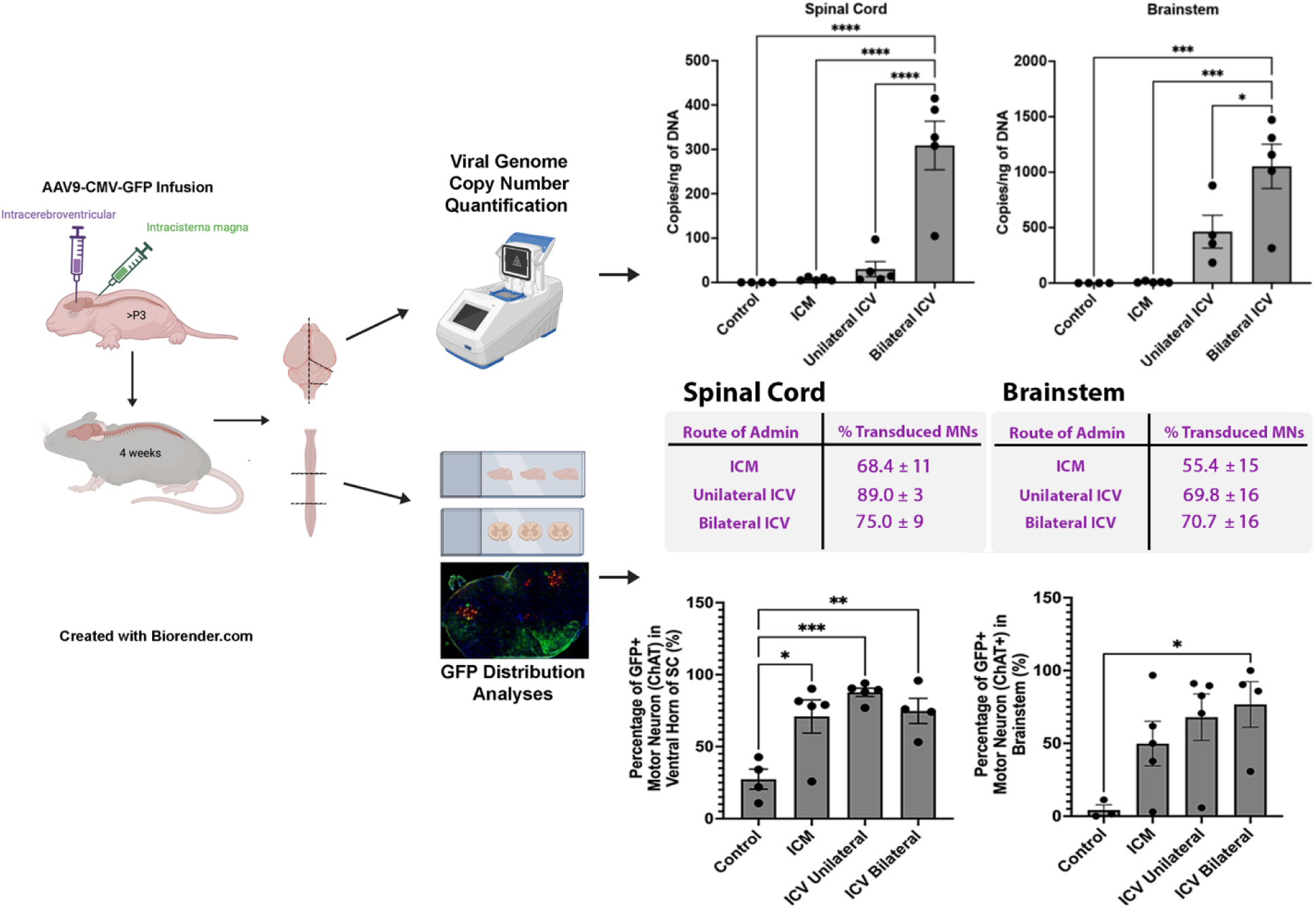

## 1 Introduction

Motor neuron diseases, such as amyotrophic lateral sclerosis (ALS) are devastating neurodegenerative conditions that are characterised by progressive degeneration of motor neurons, leading to debilitating muscle weakness, paralysis, and ultimately death. The unmet need to identify effective therapeutic strategies to target the underlying genetic and molecular abnormalities in ALS has driven the exploration of innovative approaches. The manipulation of genetic material, or gene therapy, to treat or prevent disease, has been transformative in the field of medicine.

Among various gene delivery vectors, adeno-associated virus serotype 9 (AAV9) has gained prominence as an ideal candidate for delivering therapeutic genes to the central nervous system (CNS) (Asokan et al., 2012). AAV9 has a high affinity for motor neurons, a versatile and safe profile, and has the capacity to cross the blood-brain barrier (BBB) in both rodents and non-human primates (Foust and Kaspar, 2009, Duque et al., 2009, Gray et al., 2011, Dismuke et al., 2013, Bevan et al., 2011, Fu et al., 2011). AAV9’s unique tropism towards the central nervous system has driven extensive work into its application for targeted gene delivery for neurodegenerative conditions, such as the monogenic, childhood motor neuron disease, SMA, for which gene therapies are now commercially available (Kotulska et al., 2021).

The choice of delivery method of AAV9 plays a crucial role in determining the efficiency and specificity of vector transduction within the CNS. Several strategies have been developed for AAV9 delivery targeted to the CNS including: intravenous (IV) injection; intra-cerebroventricular (ICV) injection; intrathecal (IT) injection either by lumbar puncture or intra-cisterna magna (ICM), sub-pial and intra-parenchymal injection. Each method offers unique advantages and challenges in terms of target cell specificity, transduction efficiency and off-target effects and have been reviewed recently (Peters et al., 2021). In brief, intravenous injection can provide systemic distribution of AAV9 vectors, however there are numerous challenges with systemic introduction of AAV9. Firstly, vectors are diluted in the circulation, leading to higher dosing requirements which may have toxicity and cost implications. There is also more limited BBB penetration when delivering AAV9 systemically and it may lead to heightened immune response or result in non-specific transduction in peripheral organs (Samaranch et al., 2012, Mingozzi and High, 2013). There have also been concerns raised regarding AAV-mediated genotoxicity in the liver, which remains debated in the field, and continues to be monitored in clinical trials (Chandler et al., 2015).

In contrast, direct injection into cerebrospinal fluid (CSF) allows for precise targeting of specific CNS regions or cell populations but may require more invasive procedures that pose risk of tissue damage or immune responses. Intra-cerebroventricular (ICV) and cisternal (ICM) methods offer an alternative route for delivery of AAV9 vectors directly into cerebrospinal fluid (CSF), enabling widespread distribution within the CNS, whilst minimising invasiveness and allowing for lower dosing than systemic delivery. In this study, we chose to deliver vectors early in postnatal development as this is a useful time point for therapeutic proof of concept, particularly where neurodegenerative phenotypes develop rapidly such as in models of spinal muscular atrophy. In addition, at this age, delivery is less invasive, since injections can be delivered without surgery due to skull structure. Despite extensive research on AAV9 vector delivery methods, direct comparative studies evaluating viral distribution after different administration routes in mice remain limited.

Here, we therefore aimed to compare ICV and ICM methods of AAV9 delivery to mice *in vivo* and compare motor neuron transduction levels to determine the most effective approach for specific motor neuron targeting in pre-clinical studies. We utilised AAV9 controlled cytomegalovirus (CMV)-promoter driven expression of enhanced green fluorescent protein (eGFP) to compare levels of motor neuron transduction in the spinal cord after delivery into either cisterna magna (ICM), or into the lateral ventricle (ICV) of neonatal wild-type mice. At around 4 weeks post-infusion, a timepoint at which transgene expression should be high (Stoica et al., 2013, Hollidge et al., 2022), the animal was humanely sacrificed and we used quantitative PCR and immunohistochemical approaches to assess viral distribution. We found that all CNS-directed delivery methods yielded high levels of lumbar spinal cord motor neuron transduction (>68%), however noted that upper motor neurons did not appear transduced in any case. ICM delivery was more variable and technically challenging than ICV but lead to minimal astrocytic transduction. Unilateral ICV delivery targeted lumbar motor neurons at high and consistent levels, along with astrocytes in the cortex. Bilateral ICV delivered over two days resulted in significantly higher astrocytic transduction in spinal cord, brainstem and cortex, and resulted in higher peripheral viral genome copies. Overall, we provide the first direct comparison of methods of CNS targeting using AAV9 in mice.

## 2 Methods

### 2.1 Animals

All animal procedures were performed in accordance with UK Home Office and institutional guidelines. Pregnant, wild-type C57BL/6J mice were housed in open cages in pathogen-free conditions with *ad libitum* access to food and water, on a 12:12 light/dark cycle within animal facilities at the University of Sheffield. Pregnant dams were acclimated to the smell of tattoo ink and isoflurane around one week prior to birthing to familiarise them with these scents and reduce risk of pup rejection. At birth, pups were identified by tattoos that were applied to the footpad under local anaesthetic (EMLA cream) until ear notching was possible. Injections were undertaken before postnatal day (P) 3. Both sexes were used. Runts, which were identified easily on visual inspection, were excluded from the study. At weaning, pups were group housed.

### 2.2 Injections of viral vector

Self-complimentary (sc) AAV9-CMV-eGFP vector (4815 bp, packaged size 2.2 kb) at titre 9.45E+10 viral genomes (vg)/mouse in 0.001% pluronic carrier, or vehicle only (pluronic) were delivered before postnatal day 3 under general anaesthesia (isoflurane) in the following groups, as summarised in Table 1: 5 μL intra-cisterna magna (ICM); 5 μL unilateral (right only) intra-cerebroventricular (ICV); or 5 μL unilateral (right) ICV on day 1 **and** 5 μL unilateral (left) ICV on day 2 (referred to as ICV bilateral). Controls received an equivalent injection of carrier only. Injection material was loaded into a 25 μL Hamilton syringe with 33G, small hub RN needle (PST 4).

**Table 1:**
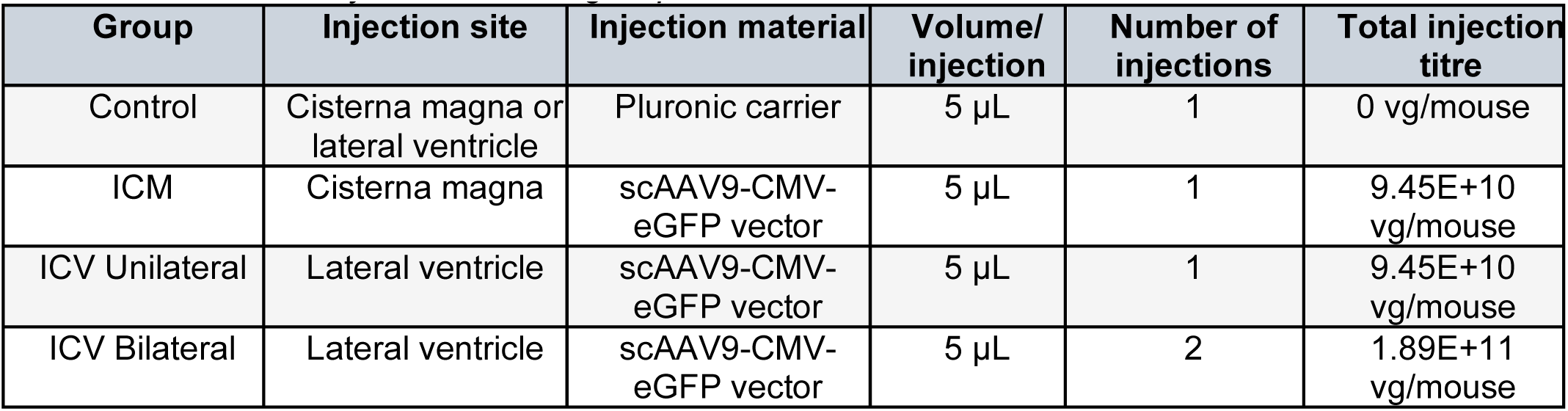
Summary of treatment groups and titres.

| Group | Injection site | Injection material | Volume/<br>injection | Number of<br>injections | Total injection<br>titre |
| --- | --- | --- | --- | --- | --- |
| Control | Cisterna magna or lateral ventricle | Pluronic carrier | 5 μL | 1 | 0 vg/mouse |
| ICM | Cisterna magna | scAAV9-CMV-eGFP vector | 5 μL | 1 | 9.45E+10<br>vg/mouse |
| ICV Unilateral | Lateral ventricle | scAAV9-CMV-eGFP vector | 5 μL | 1 | 9.45E+10<br>vg/mouse |
| ICV Bilateral | Lateral ventricle | scAAV9-CMV-eGFP vector | 5 μL | 2 | 1.89E+11<br>vg/mouse |

Anaesthesia was induced at 5% isoflurane, and 3 L O^2^/min. Pups were transferred onto a Model 940 Small Animal Stereotaxic Instrument (Bilaney) and placed over a WeeSight™ transilluminator to visualise the injection site. Pups were maintained under anaesthesia on a modified nosecone at 2% isoflurane, 0.5 L O^2^/min throughout, with alterations based on individual monitoring. For ICM, the injection site was visible through the skin as a small dark triangle and was most easily accessed when the pups head was tilted slightly forward over the transilluminator. Infusions were initiated immediately after the needle punctures through the skin. For ICV, the site was located approximately 0.25 mm lateral to the sagittal suture and 0.50 - 0.75 mm rostral to the neonatal coronary suture and at a depth of 2 mm, as described previously by Glascock et al., 2011 (Glascock et al., 2011). 5 μL volumes were delivered by syringe driver (Harvard Apparatus) at a rate of 1 μL/minute. Note that the needle was left in position for around 1 minute after infusion completion to allow pressure to dissipate and then the needle was withdrawn slowly to minimise flowback. After infusion, pups were recovered upright (in a tube holder) to encourage spinal distribution, in an incubator, for around 10 minutes before returning to the home cage and monitoring for resumption of normal maternal care.

### 2.3 Tissue Collection

Around 4 weeks post-injection, mice were sacrificed by an overdose of pentobarbitol and a cardiac perfusion was performed with PBS. The spinal cord was then dissected *in situ*; the start of the lumbar region was identified anatomically by the end of the ribcage and presence of the lumbar enlargement. The lumbar region was fixed in 4% paraformaldehyde overnight. The remaining spinal cord was snap frozen on dry ice for viral copy number analysis. The brain was dissected and split along the midline, with the right half being fixed in 4% paraformaldehyde overnight, and left being split into regions of cortex, cerebellum and brainstem for copy number analysis. Liver and heart were also snap frozen to assess off-target effects.

### 2.4 Immunohistochemistry

Fixed brains and spinal cords were paraffin embedded and sagittally or coronally sectioned on a microtome at 10 μm respectively. Tissue was deparaffinised and subjected to antigen retrieval in a pressure cooker (Access Revelation, pH 6.4). Sections were blocked with 5% BSA and 0.25% tx-100. Slides were then immunostained for: GFP + ChAT (motor neuron identity); GFP + GFAP (astrocytic identity); and GFP + Iba1 (microglial identity). Note that for double stains using ChAT, we found that staining was optimal when antibodies were added consecutively. Primary and secondary GFP staining was therefore performed and then followed by ChAT staining. For all other combinations, primary antibodies were added simultaneously in a single incubation. A summary of primary antibodies and conditions of use are shown in Table 2. Secondary antibodies used included: goat anti-chicken Alexa Fluor 488, 1:1000 (ThermoFisher, A-11039); donkey anti-chicken Alexa Fluor 488, 1:1000 (Jackson Immunoresearch, 703-545-155); goat anti-rabbit Alexa Fluor 555, 1:1000 (ThermoFisher, A27039); donkey anti-goat Alexa Fluor 555, 1:400 (Thermofisher A21432). Negative controls consisted of sections incubated in the absence of primary antibody.

**Table 2:**
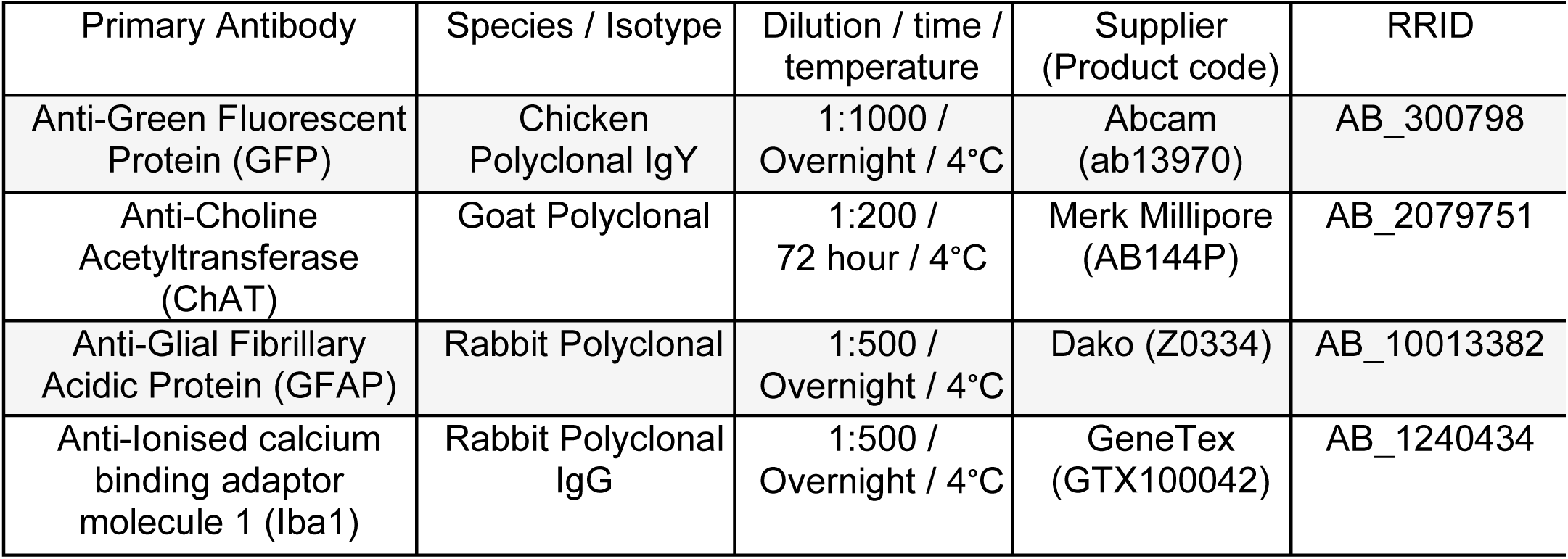
Summary of primary antibodies used for immunohistochemistry.

### 2.5 Viral DNA extraction

DNA was extracted from cortex, cerebellum, brainstem, spinal cord, liver and heart by standard phenol-chloroform extraction method. In brief, tissue was crushed with a pestle in a 1.5mL Eppendorf, and 250 μL of reporter lysis buffer (Promega) was added and tissue was crushed again. 1 μL of 20 mg/mL proteinase K was added, pulse vortexed and incubated at 50°C for 1 hour. Samples were homogenised 15x with a 21G needle. 250 μL of phenol/chloroform/isoamyl alcohol was added and vortexed vigorously for 3 minutes. Samples were centrifuged at 17000 x g at room temperature for 10 minutes. Around 225 μL of supernatant was removed to a new 1.5 mL Eppendorf and 25μL 3M sodium acetate and 1 μL glycogen was added and pulse vortex. 750 μL of 100% ethanol was added and tubes were inverted. Samples were left to precipitate overnight at −20ᵒC. The following day, samples were centrifuged at 17000 x g at room temperature for 20 minutes. Supernatant was discarded and samples was washed with 500 μL of 70% ethanol and centrifuged at 17000 x g at room temperature for 10 minutes. Supernatant was discarded, and the sample was pulse spun and a pipette was used to remove as much ethanol as possible. Samples were left to air dry for at least an hour at room temperature. DNA was resuspended in 25 - 50 μL of RNAse free water. Concentration and quality were determined by nanodrop. DNA was stored at −20°C.

### 2.6 Quantitative Polymerase Chain Reaction (qPCR)

A BioRad CFX Touch Real-Time PCR qPCR system was used to determine the number of viral genome copies present in different tissues. Primers were used that targeted the bovine growth hormone (bGH) polyA tail of the pAAV9-CMV-eGFP vector: bGH pA Forward CTCGACTGTGCCTTCTAGTTG and bGH pA Reverse CCTACTCAGACAATGCGATG. A standard plasmid (CMV.GFP plasmid clone 675.5, Vector Biolabs) was used to generate a standard reference curve with a known number of viral copies (see Appendix, Fig. A. 1 for curve). SYBR green was used with cycling times as shown in Table 3.

**Table 3:**
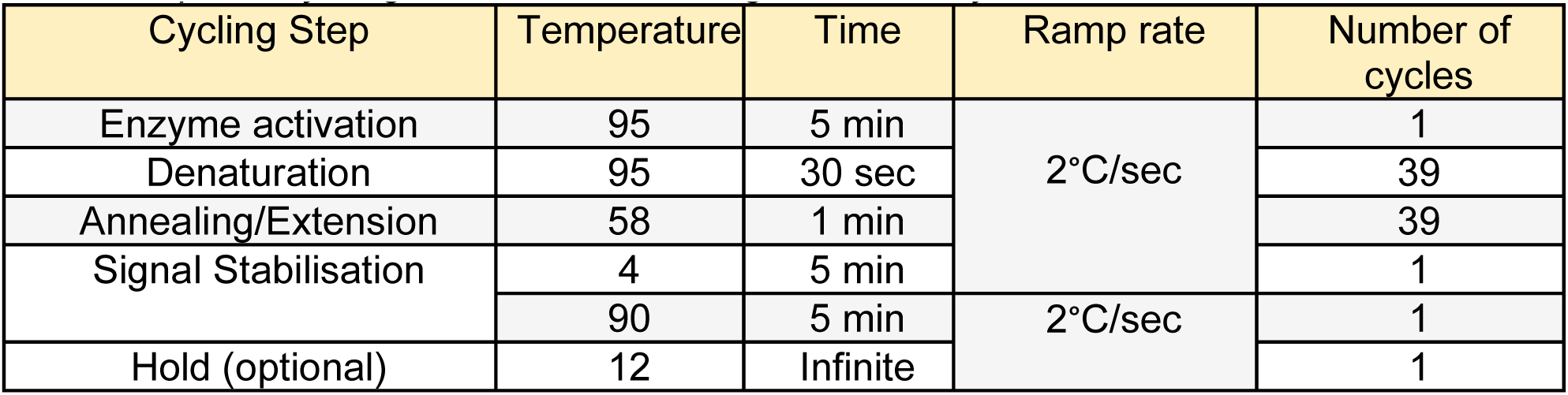
qPCR cycling conditions for viral genome analysis.

### 2.7 Quantification and Statistical Analysis

Images of spinal cord and brainstem sections were taken on an Incell analyser 2200/2000 with equal exposure settings, before being taken into Image J and adjusted for brightness and contrast across all sections, and a flat-field correction was applied. A minimum of 3 slides per animal were used for quantification, with a minimum of 65 motor neurons per animal. Motor neurons were manually quantified for co-localisation of anti-ChAT, indicative of motor neuron identity, and anti-GFP, with the criteria that motor neurons must also have a positive DAPI nuclei. For cortical anti-GFAP quantification, indicative of astrocytes, and anti-GFP quantification, Image J was used to count cells and co-localisation assessed on overlap. A manual count was then taken of GFP positive cells, to allow a correction to be applied for non-selective staining of blood vessels. Quantification of anti-ChAT and anti-GFAP, were analysed using CellProfiler^TM^ (software available www.cellprofiler.org and pipelines available https://github.com/CFSander/CellProfiler_astrocyte_pipelines.git) (See Fig. A. 2 for example training paradigm) (Lamprecht et al., 2007. Stirling et al., 2021). Investigators were blinded to treatment group during all manual quantification, however blinding was not necessary for automated pipelines. Data were collated in excel and statistical analyses were undertaken in Prism10.

## 3 Results

### 3.1 AAV9 delivery to CSF leads to high motor neuron transduction in the lumbar spinal cord, with unilateral ICV delivery providing consistently high levels

There is a distinct lack of literature that directly compares transduction levels *in vivo* in mice following different CNS-directed delivery methods of AAV9. We therefore firstly aimed to quantify the number of transduced motor neurons in the spinal cord 4 weeks after ICM, unilateral ICV or bilateral ICV delivery of AAV9-CMV-eGFP.

We used immunohistochemical approaches (**Figure 1**) to manually count the number of motor neurons that co-localised with choline acetyltransferase (ChAT), indicating motor neuron identity, and expressed this as a percentage of the total number of motor neurons counted in the lumbar spinal cord (**Figure 2**). Note that, unexpectedly, we found cells that were quantified as positive for GFP in control animals. We are confident that this positivity is artefact, since viral genome copy number analysis in control animals was also negligible (see Table 6). We acknowledge, however, that it is possible that we may have overestimated percentage transduction in injected animals due to this background artefact. Note that all quantifications were either automated or performed blinded. For transparency, we have included control data for reference and to highlight the challenges with eliminating all background staining with GFP immunohistochemistry.

**Figure 1:**
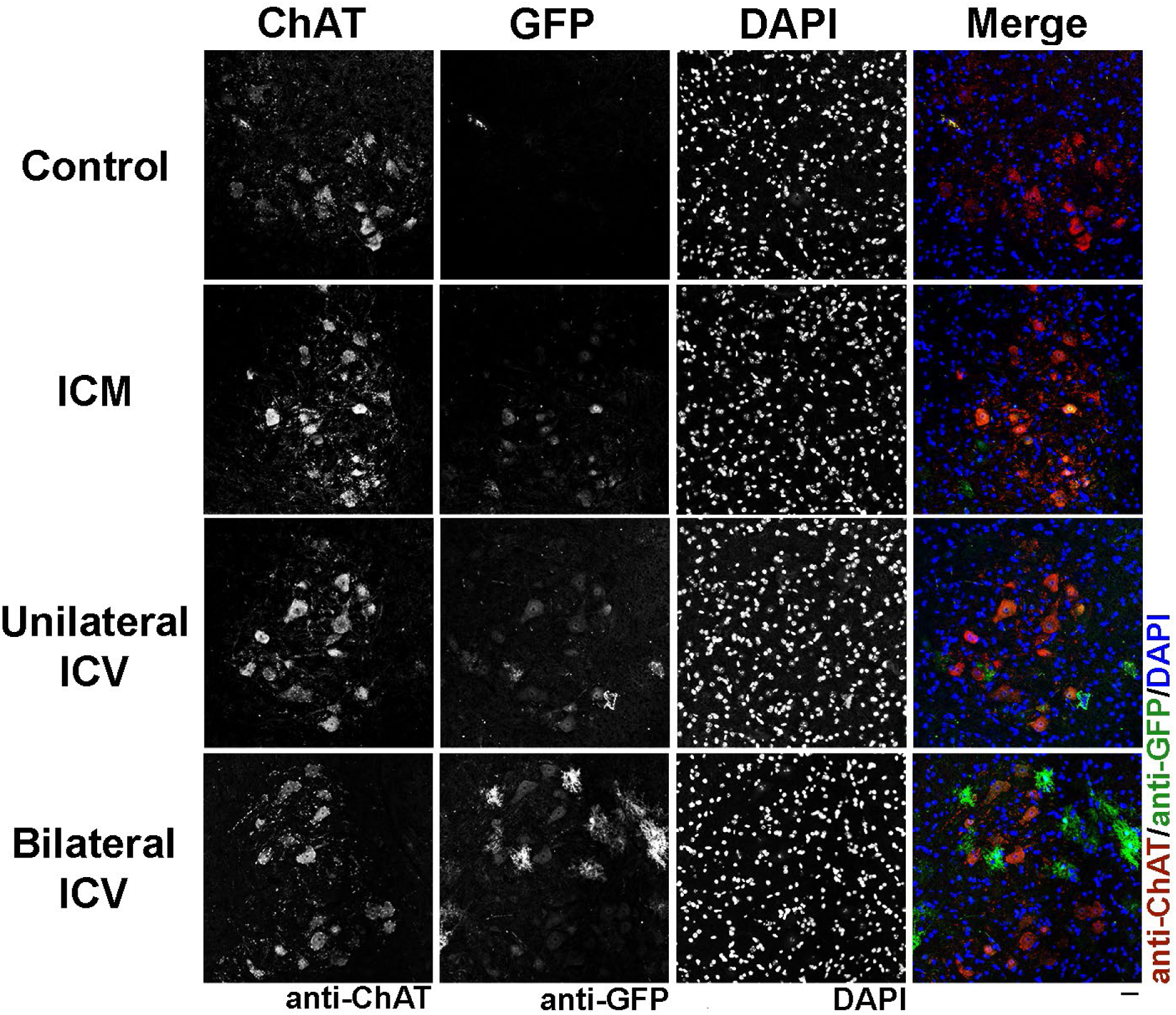
Visualisation of choline acetyltransferase (ChAT) positive cells, indicating motor neurons, and GFP co-localisation in the ventral horn of the spinal cord in mice around 4 weeks after AAV9-CMV-eGFP delivery. Note that transduction can be clearly visualised in all delivery conditions. Note that there are transduced cell types particularly visible in the bilateral ICV condition which are not motor neurons and which we later identified as astrocytes. Sections labelled with anti-ChAT, anti-GFP and DAPI. Scale bar = 40 µm.

**Figure 2:**
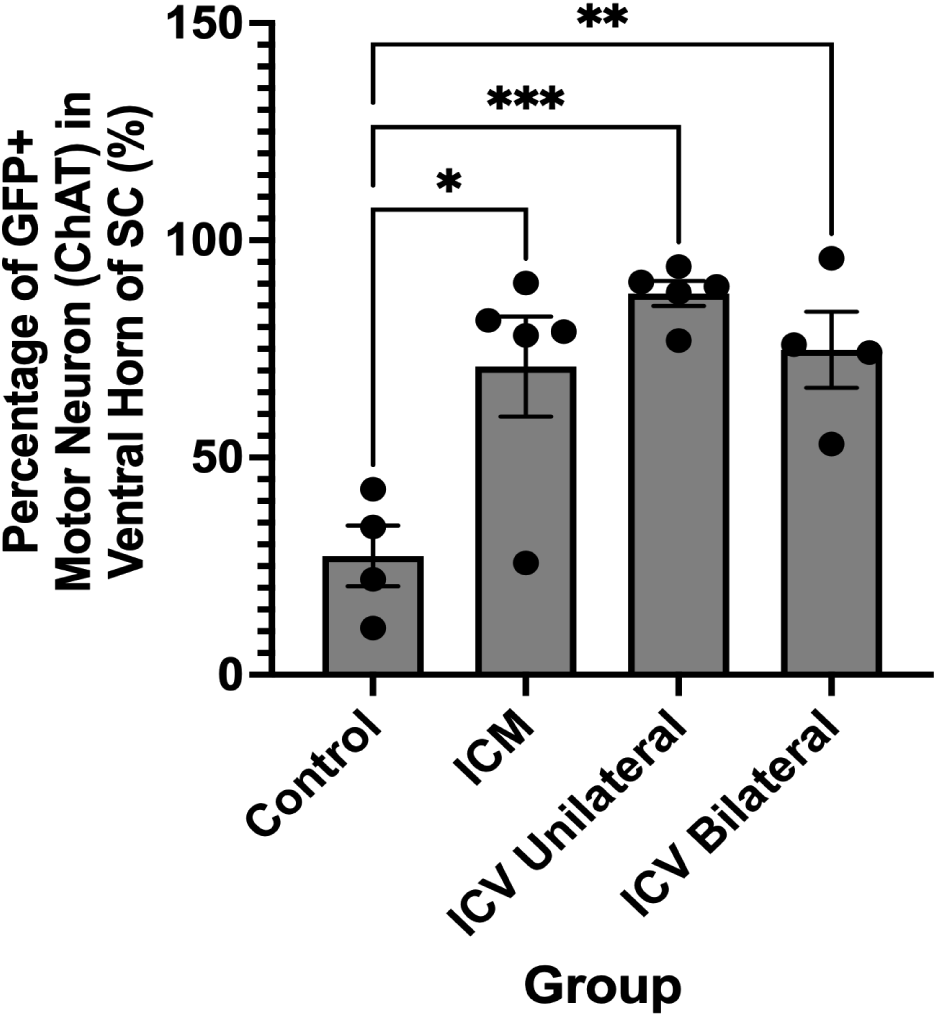
Quantification of the percentage of GFP positive cells that are co-localised with ChAT, indicating transduced motor neurons, in the spinal cord after different CNS-directed delivery methods of AAV9-CMV-eGFP. Note that there are high levels of transduction in all groups, however the most consistent and highest transduction is seen after unilateral ICV delivery. ICM = intra-cisterna magna; ICV unilateral = unilateral intra-cerebroventricular; ICV bilateral = bilateral intra-cerebroventricular delivery. One-way ANOVA with Tukey’s correction for multiple comparisons. *p≤0.05, **p≤0.01 ***p≤0.001. Error bars = mean ± SEM.

We found that all delivery methods lead to high levels of motor neuron transduction in the spinal cord, with over 68% of lumbar motor neurons being transduced in all groups, as summarised in Table 4. We noted that unilateral ICV approaches yielded more consistent, higher percentages of transduced motor neurons in the lumbar spinal cord than other approaches (**Figure 2**).

**Table 4:** MN counts in lumbar spinal cord and average transduction level.

| Route of Administration | Total MNs counted (ChAT+) | Transduced MNs (eGFP+/ChAT+) | Percentage of Transduced MNs ±SEM (%) |
| --- | --- | --- | --- |
| ICM (n=5) | 661 | 452 | 68.4 ± 11 |
| Unilateral ICV (n=5) | 744 | 662 | 89.0 ± 3 |
| Bilateral ICV (n=4) | 643 | 482 | 75.0 ± 9 |

### 3.2 AAV9 delivery to CSF leads to high motor neuron transduction in the brainstem but not in the motor cortex

We next sought to investigate and compare the extent to which motor neurons in the brainstem and cortex were targeted by ICM and ICV delivery methods of AAV9-CMV-eGFP.

Using immunohistochemistry (**Figure 3**), we manually quantified the co-localisation between ChAT, indicative of motor neurons, and GFP, indicating positive transduction in the brainstem and expressed the number of co-localised motor neurons as a percentage of total motor neurons counted (**Figure 4**).

**Figure 3:**
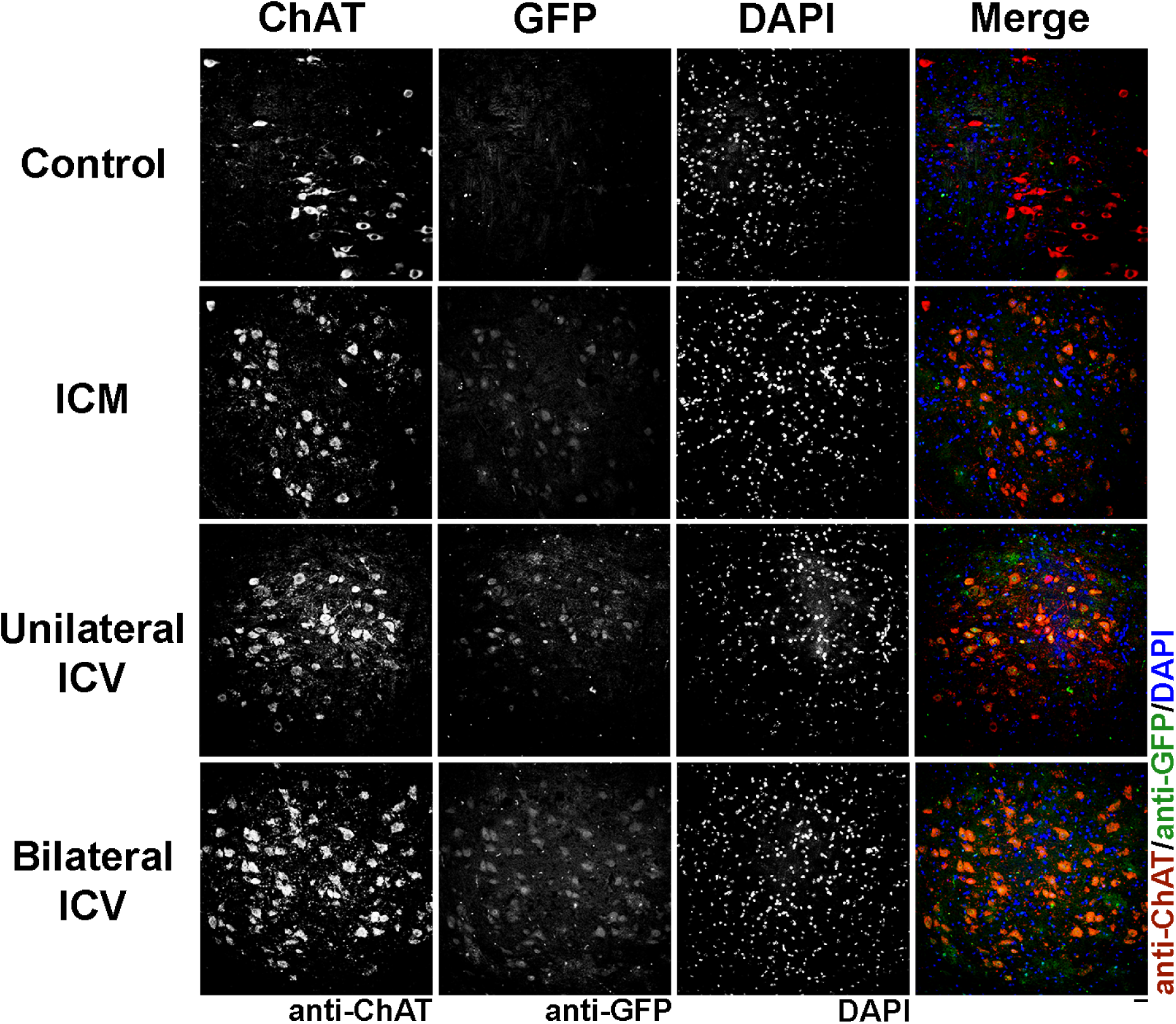
Visualisation of choline acetyltransferase (ChAT) positive cells, indicating motor neurons, and GFP co-localisation in the brainstem in mice around 4 weeks after AAV9-CMV-eGFP delivery. Note that transduction can be clearly seen in all delivery conditions. Sections are labelled with anti-ChAT, anti-GFP and DAPI. Scale bar = 20 µm.

**Figure 4:**
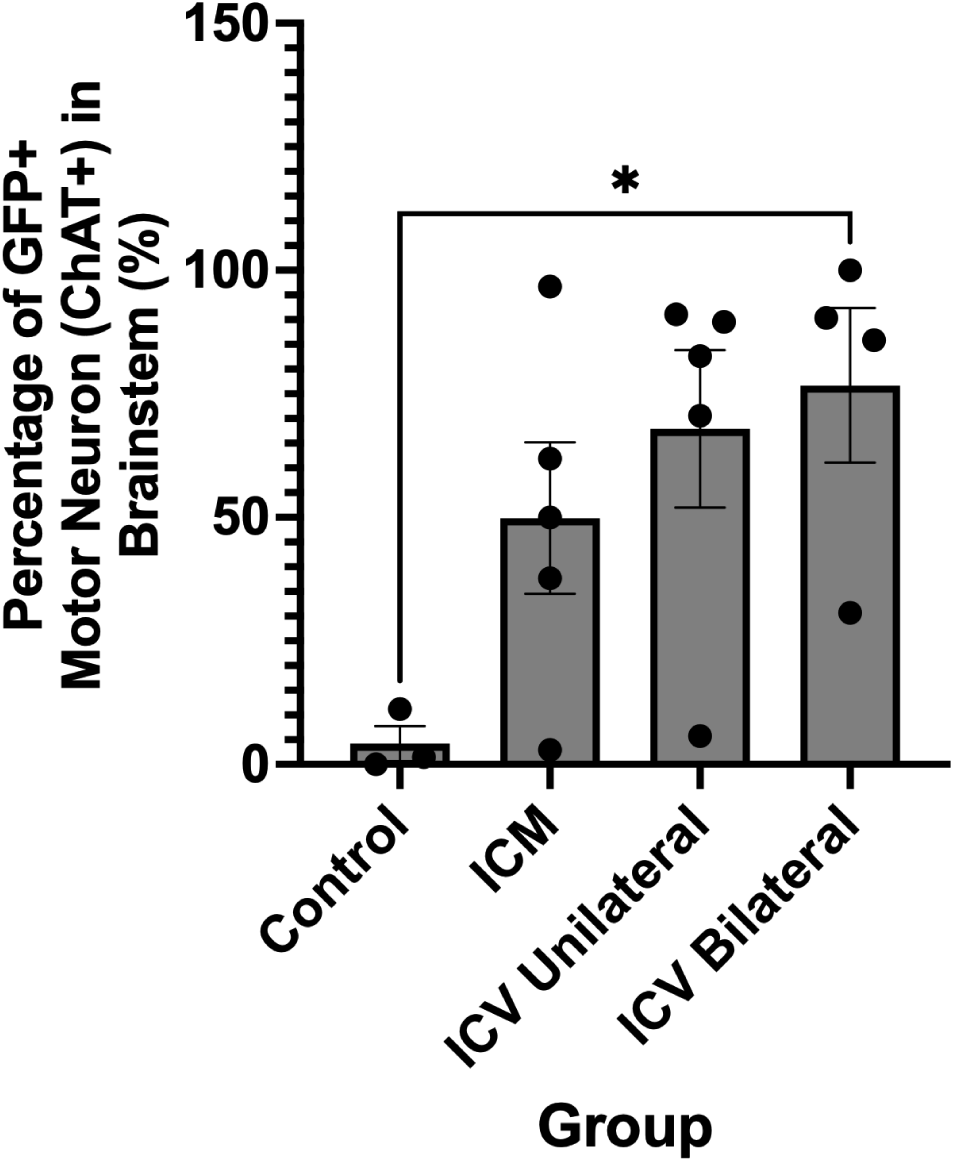
Quantification of the percentage of GFP positive cells that are co-localised with ChAT, indicating transduced motor neurons, in the brainstem after different CNS-directed delivery methods of AAV9-CMV-eGFP. Note that there are high levels of transduction in all groups, although this is variable between animals. ICM = intra-cisterna magna; ICV unilateral = unilateral intra-cerebroventricular; ICV bilateral = bilateral intra-cerebroventricular delivery. One-way ANOVA with Tukey’s correction for multiple comparisons. *p≤0.05. Error bars = mean ± SEM.

We found that motor neuron transduction was high in the brainstem in both conditions, with >55% of motor neurons transduced in all groups, as summarised in Table 5, however we noted high variability (**Figure 4**).

**Table 5:** MN counts in brainstem and average transduction level.

| Route of Administration | Total MNs counted (ChAT+) | Transduced MNs (eGFP+/ChAT+) | Percentage of Transduced MNs ± SEM (%) |
| --- | --- | --- | --- |
| ICM (n=5) | 473 | 262 | 55.4 ± 15 |
| Unilateral ICV (n=4) | 424 | 296 | 69.8 ± 16 |
| Bilateral ICV (n=5) | 850 | 601 | 70.7 ± 16 |

**Table 6:** Viral copy number analysis in central tissues as determined by qPCR.

| Tissue | Number of viral copies/ng of DNA (Mean ± SEM) | | | |
| --- | --- | --- | --- | --- |
| | Control (n=4) | ICM (n=5) | Unilateral ICV (n≥3) | Bilateral ICV (n≥4) |
| Spinal cord | 0 | 7.7 ± 2 | 30.1 ± 17 | 308.9 ± 55 |
| Brainstem | 0 | 9.1 ± 3 | 463.5 ± 148 | 1053.0 ± 200 |
| Cerebellum | 0 | 65.4 ± 40 | 6.9 ± 3 | 46.5 ± 17 |
| Cortex | 0 | 56.9 ± 54 | 5598.0 ± 3688 | 3432.0 ± 1101 |

In the cortex, however, we were unable to detect co-localisation of the neuronal marker (anti-NeuN) with GFP, suggesting minimal neuronal transduction (**Figure 5**).

**Figure 5:**
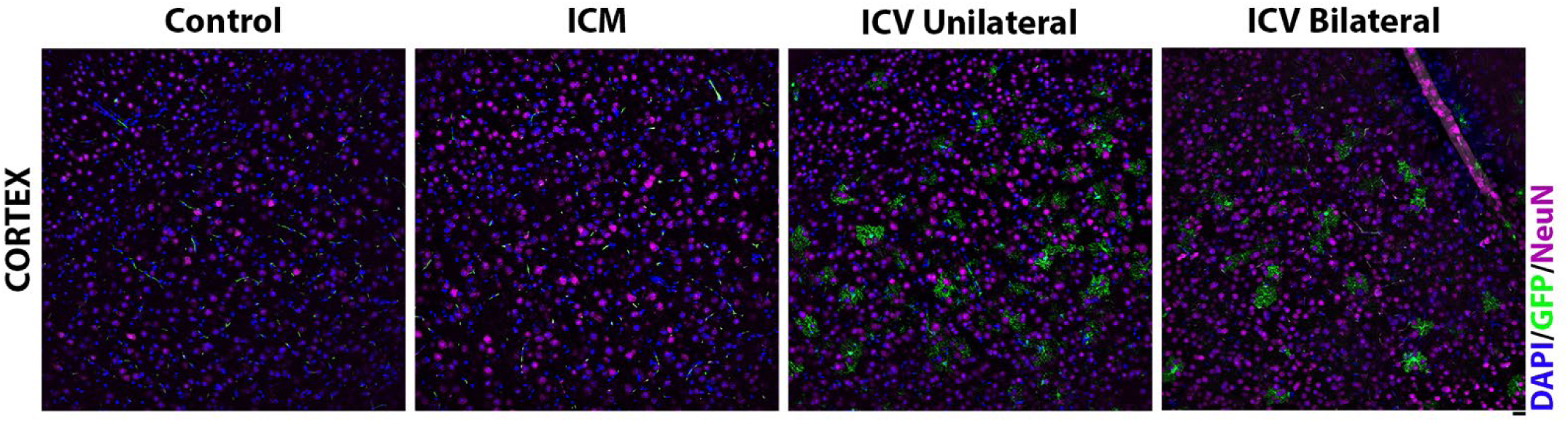
Visualisation of the neuronal marker, anti-NeuN, and GFP in the motor cortex in mice around 4 weeks after AAV9-CMV-eGFP delivery. Note that a lack of co-localisation is seen with NeuN, suggesting minimal neuronal transduction. Note that we can see other non-neuronal cell types that are transduced following ICV administration in the cortex. Scale = 20*µ*m.

### 1.1 The number of viral genome copies in spinal cord and brainstem is higher with ICV delivery methods

Quantification of viral genome copy number can provide additional insight into the biodistribution and transduction efficiency of gene therapy vectors. Here, we used quantitative PCR (qPCR) on genomic DNA isolated from various CNS tissues to gain further insight into viral distribution and compare this across delivery methods.

We quantified the average number of viral copies/ng of DNA in each tissue (Table 6 and Figure 6) by extrapolating Ct values from a standard curve generated from a GFP plasmid with known viral copy number (see Supplementary data, Fig.A1). As expected, we found that the control group had negligible viral copy numbers, confirming that our detection method was specific, and further evidencing that immunohistochemical positivity previously detected in control is artefact.

**Figure 6:**
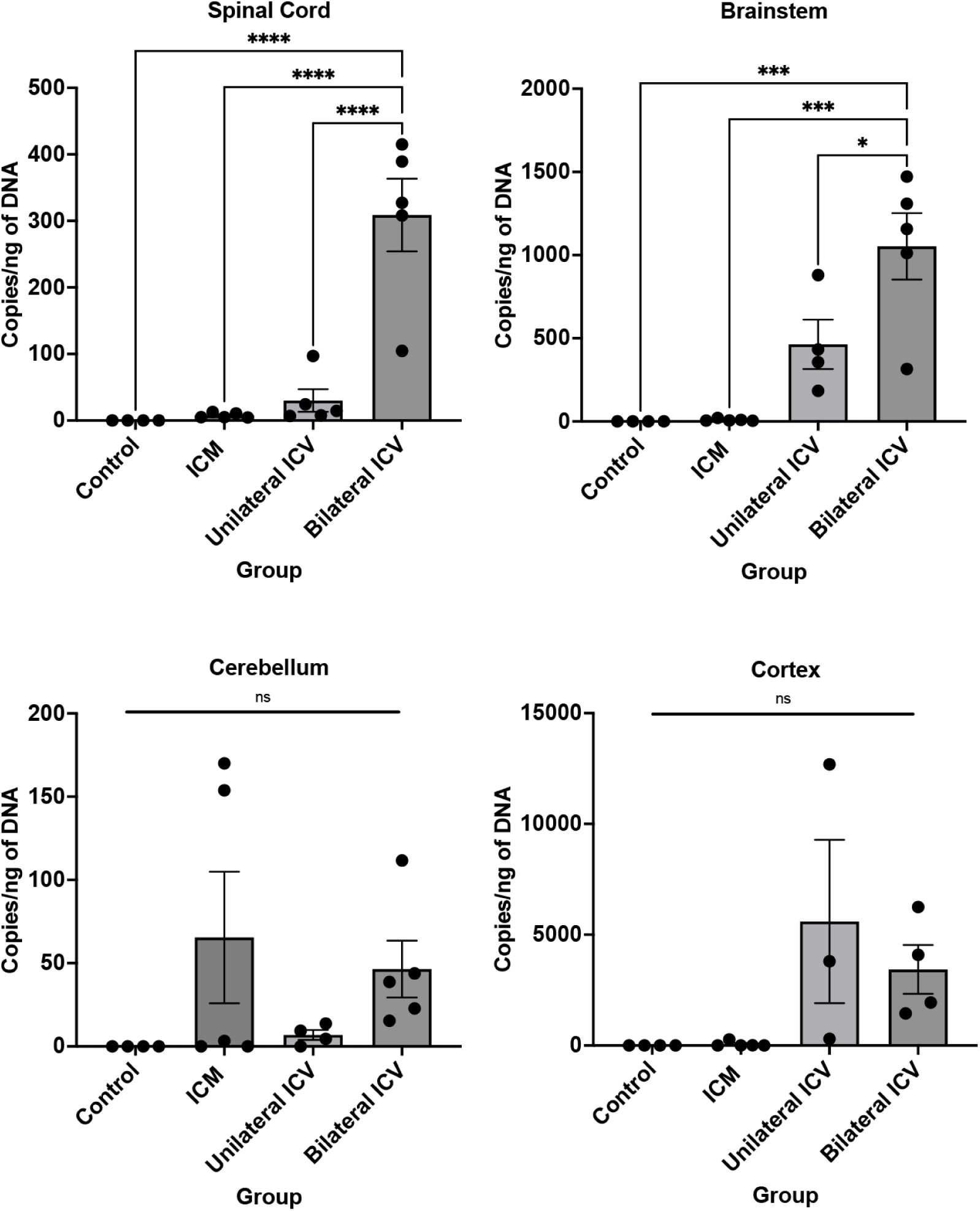
Quantification of viral copy number in spinal cord, brainstem, cerebellum and cortex. Quantitative PCR on CNS tissue suggests that there are high copy numbers in spinal cord and brainstem, with a trend towards higher copy number in these tissues when delivered intra-cerebroventricularly (ICV). ICM = intra-cisterna magna; ICV unilateral = unilateral intra-cerebroventricular; ICV bilateral = bilateral intra-cerebroventricular delivery. One-way ANOVA with Tukey’s correction for multiple comparisons. ns=not significant, *p≤0.05, ***p≤0.001, ****p≤0.0001. Error bars = mean ± SEM.

We were able to detect viral copies in all treatment groups. There was a trend towards a higher viral copy number in spinal cord and brainstem after ICV delivery methods.

In spinal cord, there were between 7-8 copies/ng of DNA after ICM delivery, threefold higher following unilateral ICV at around 30 copies/ng of DNA and significantly higher after bilateral ICV at around 308 copies/ng of DNA, as summarised in Table 6 and Figure 6. The viral genome copy number does not necessarily align with data obtained for motor neuron transduction rates obtained from immunohistochemical analysis (Figures 2 and 4). We can attribute this to either transduction of other cell types, such as astrocytes, or an upper limit on transduction efficiency such that higher viral genome copy number does not translate to a greater transduction rate.

A similar trend was also seen in brainstem, with around 9 copies/ng of DNA after ICM, 464 copies/ng of DNA after ICV unilateral and 1053 copies/ng of DNA after bilateral ICV, also summarised in Table 6 and Figure 6. We noted increased variability in viral copy number at higher levels.

In cortex, we observed high viral genome copy numbers following ICV delivery, despite a lack of neuronal transduction. Taking copy number results together with immunohistochemical assessment, this is likely due to the high level of astrocyte transduction in cortex (see Results, Table 7 and Fig. 7).

**Figure 7:**
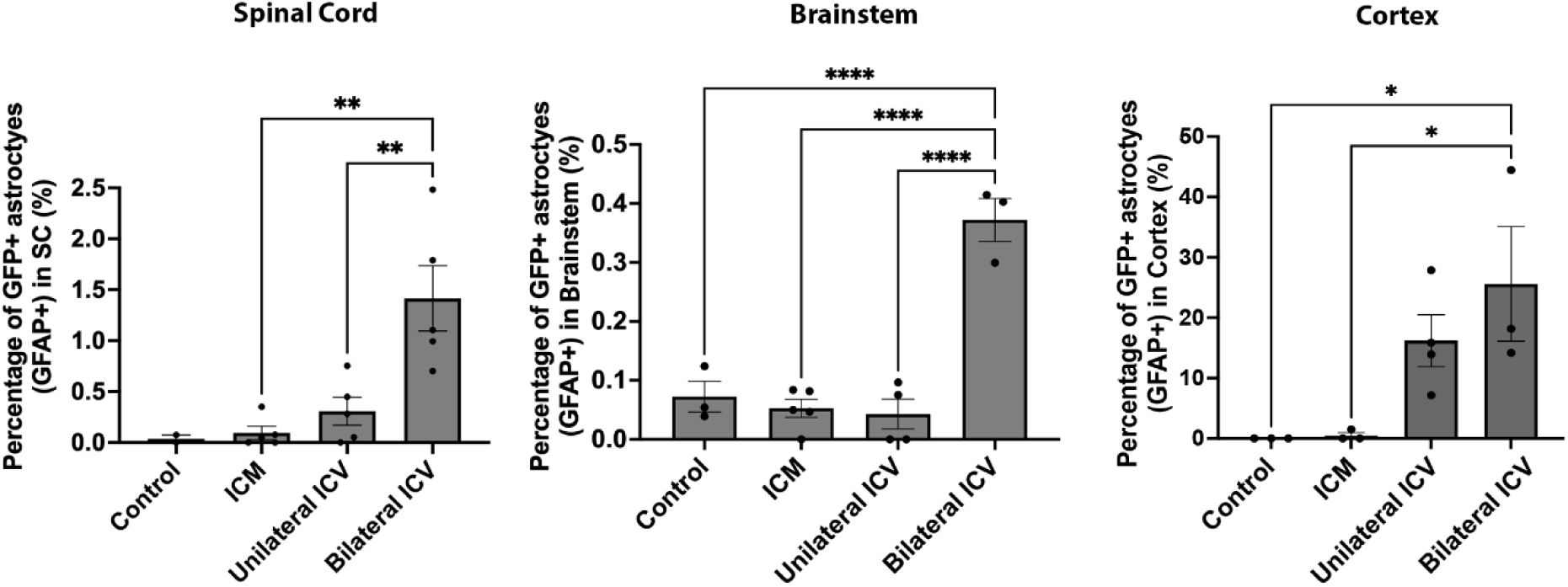
Quantification of the percentage of transduced astrocytes in spinal cord, brainstem and cortex. Quantification of GFP positive cells that co-localised with the astrocytic marker, GFAP, showed that percentages of transduced astrocytes following intra-cisternal magna (ICM) and unilateral intra-cerebroventricular (ICV) injection are not significantly different than control, suggesting minimal astrocytic transduction in these groups. Note that there are a small but significantly higher percentage of transduced astrocytes following bilateral ICV delivery in spinal cord and brainstem. Note that in cortex, ICV delivery appears to target a much higher percentage of astrocytes, compared to ICM. One-way ANOVA with Tukey’s correction for multiple comparisons. *p≤0.05, **p≤0.01, **** p≤0.0001 Error bars = mean ± SEM.

**Table 7:** Percentage of transduced astrocytes in spinal cord, brainstem and cortex.

| Tissue | Percentage of transduced astrocytes (Mean ± SEM) | | | |
| --- | --- | --- | --- | --- |
| | Control (n≥2) | ICM (n≥3) | Unilateral ICV (n≥4) | Bilateral ICV (n≥3) |
| Spinal Cord | 0.04 | 0.10 ± 0.07 | 0.31 ± 0.14 | 1.42 ± 0.32 |
| Brainstem | 0.07 ± 0.03 | 0.05 ± 0.02 | 0.04 ± 0.03 | 0.37 ± 0.04 |
| Cortex | 0 ± 0 | 0.50 ± 0.50 | 16.20 ± 4.31 | 25.60 ± 9.50 |

### 1.2 ICV delivery of AAV9-CMV-eGFP results in significantly higher astrocytic transduction than other delivery methods, particularly in cortex

In this work, we intended to selectively target motor neurons, however, it was evident from immunohistochemistry that other cell types had been transduced after AAV9-CMV-eGFP delivery that were not ChAT- or NeuN-positive. Using immunohistochemistry, we identified these cells as being positive for the astrocytic marker, glial fibrillary acidic protein (GFAP) in spinal cord, brainstem and cortex (**Figure 8**, **Figure 9** and **Figure 10** respectively). These cells also appeared negative for the microglial marker, anti-ionised calcium binding adaptor molecule 1, Iba1 (**Figure 11**), indicating that these cells were likely astrocytic in identity.

**Figure 8:**
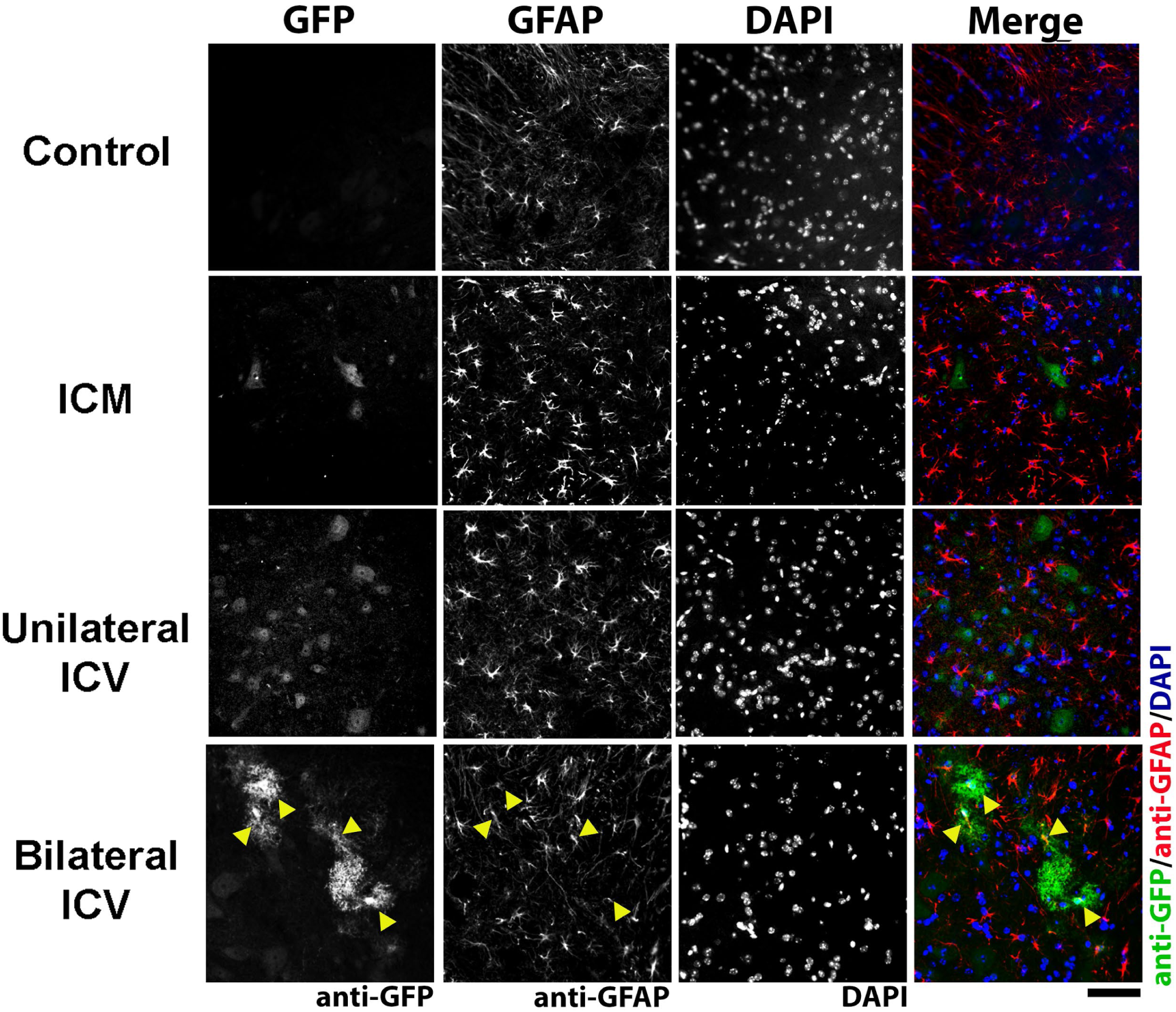
Visualisation of glial fibrillary acidic protein (GFAP) positive cells, indicating astrocytes, and GFP co-localisation in the spinal cord in mice around 4 weeks post AAV9-CMV-eGFP delivery. Note that transduction of astrocytes can be clearly seen in bilateral ICV conditions (examples indicated by yellow arrowheads). Sections are labelled with anti-GFP, anti-GFAP and DAPI. Scale bar = 50 µm.

**Figure 9:**
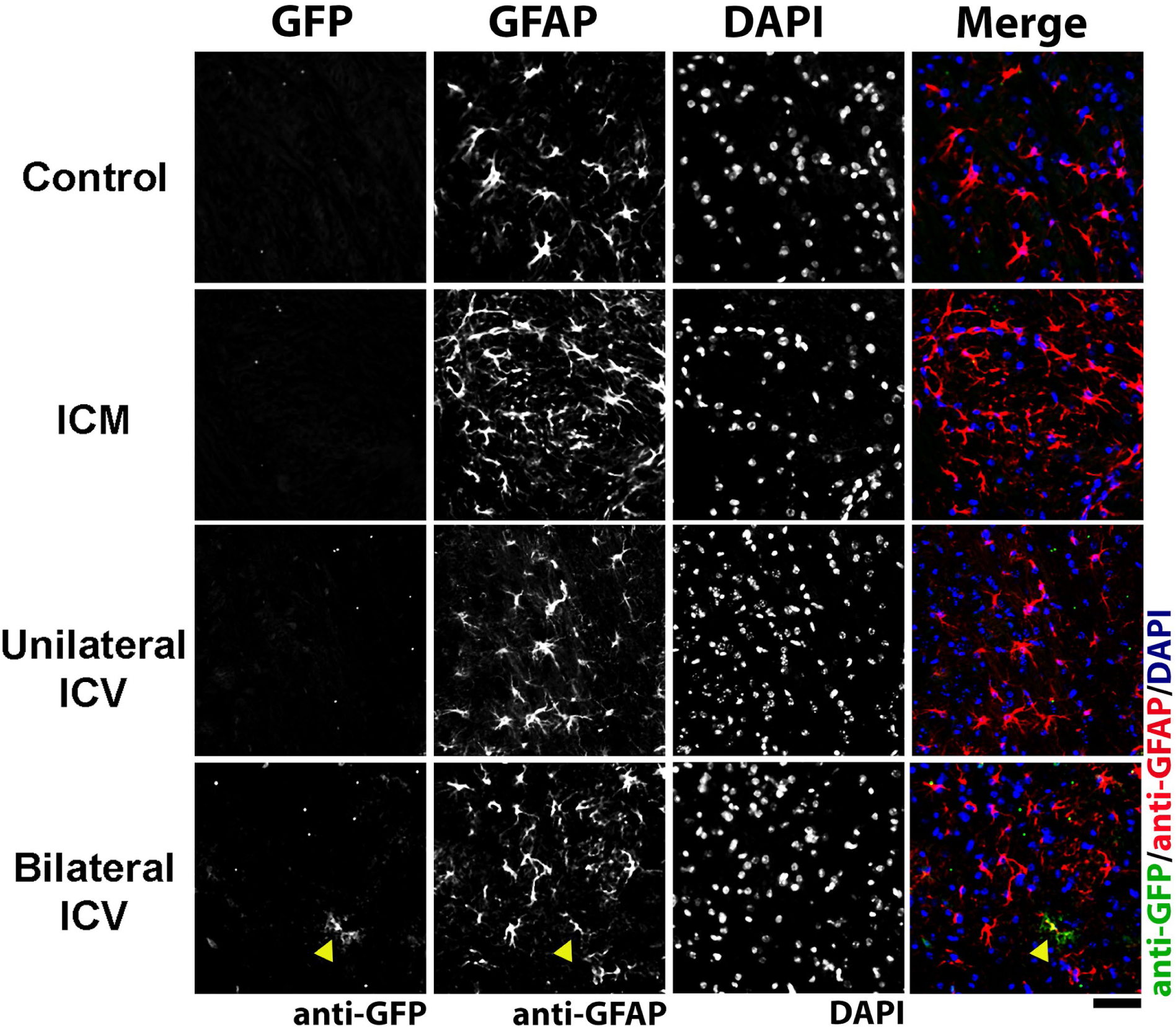
Visualisation of glial fibrillary acidic protein (GFAP) positive cells, indicating astrocytes, and GFP co-localisation in the brainstem in mice around 4 weeks post AAV9-CMV-eGFP delivery. Note that transduction of astrocytes can be clearly seen in bilateral ICV group (example indicated by yellow arrowhead). Sections are labelled with anti-GFP, anti-GFAP and DAPI. Scale bar = 50 µm.

**Figure 10:**
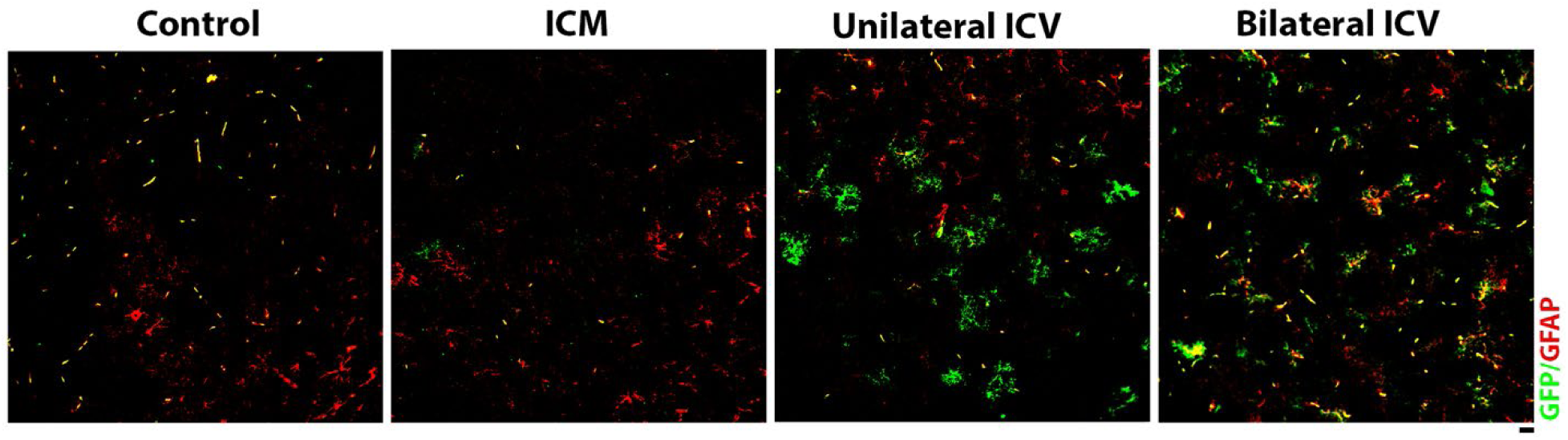
Visualisation of glial fibrillary acidic protein (GFAP) positive cells, indicating astrocytes, and GFP co-localisation in the cortex in mice around 4 weeks post AAV9-CMV-eGFP delivery. Note that transduction of astrocytes can be clearly seen in both ICV groups. Sections are labelled with anti-GFP, anti-GFAP and DAPI. Scale bar = 20 µm.

**Figure 11:**
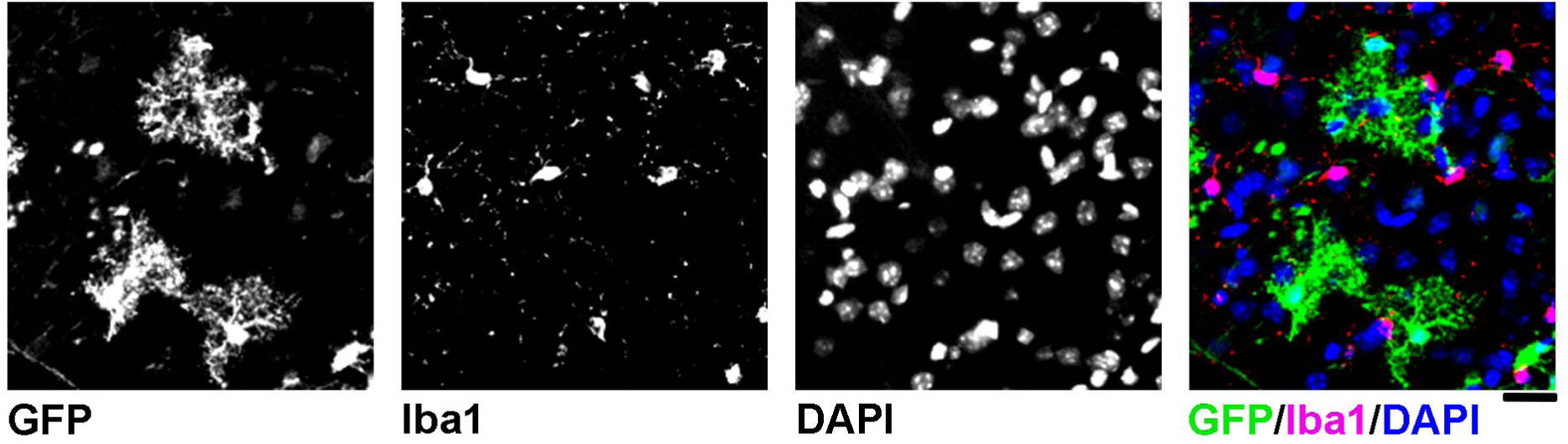
Micrograph showing a lack of GFP co-localisation with the microglial marker, anti-ionised calcium binding adaptor molecule 1, Iba1. Sections are labelled with anti-GFP, anti-Iba1 and DAPI. Scale bar = 20 µm.

We quantified the number of GFP positive cells that co-localised with GFAP and expressed this as a percentage of the total number of GFAP positive cells. As summarised in Table 7, we found that there were significantly higher percentages of astrocytes transduced with Bilateral ICV approaches in spinal cord and, particularly in the cortex. Astrocytic transduction was minimal with ICM delivery in all regions. In the ICV groups, we found significantly more astrocytic transduction in spinal cord and brainstem.

In the bilateral ICV group, in spinal cord, we found around 1.4% of astrocytes were transduced, and in brainstem 0.37% of astrocytes were transduced (Table 7, **Figure 7**).

Overall, we found that ICV delivery resulted in significantly higher astrocytic transduction than ICM delivery, although it is worth noting that numbers were low in spinal cord and brainstem, but high in cortex (Table 7, **Figure 7**). Further work is required to determine the biological significance of this, and to assess routes for better targeting of upper motor neurons.

### 1.3 Unilateral ICV delivery of AAV9-CMV-eGFP provides the lowest levels of peripheral effects

With AAV-mediated gene therapies, there is concern regarding peripheral, off-target effects, particularly pertaining to hepatotoxicity. We therefore quantified the number of viral genome copies found in liver and heart, to quantify the peripheral viral load following different CNS-directed delivery methods.

As summarised in Table 8 and Figure 12, after ICM administration there were around 9 and 14 copies in the liver and heart respectively. Unilateral ICV delivery resulted in the lowest and most consistent number of copies, with <3 copies/ng of DNA in liver and heart. As might be expected with a higher dose, bilateral ICV resulted in significantly more viral copies than any other delivery route, with around 35 and 23 copies/ng of DNA in liver and heart respectively, although we did not see any overt signs of toxicity in study animals.

**Figure 12:**
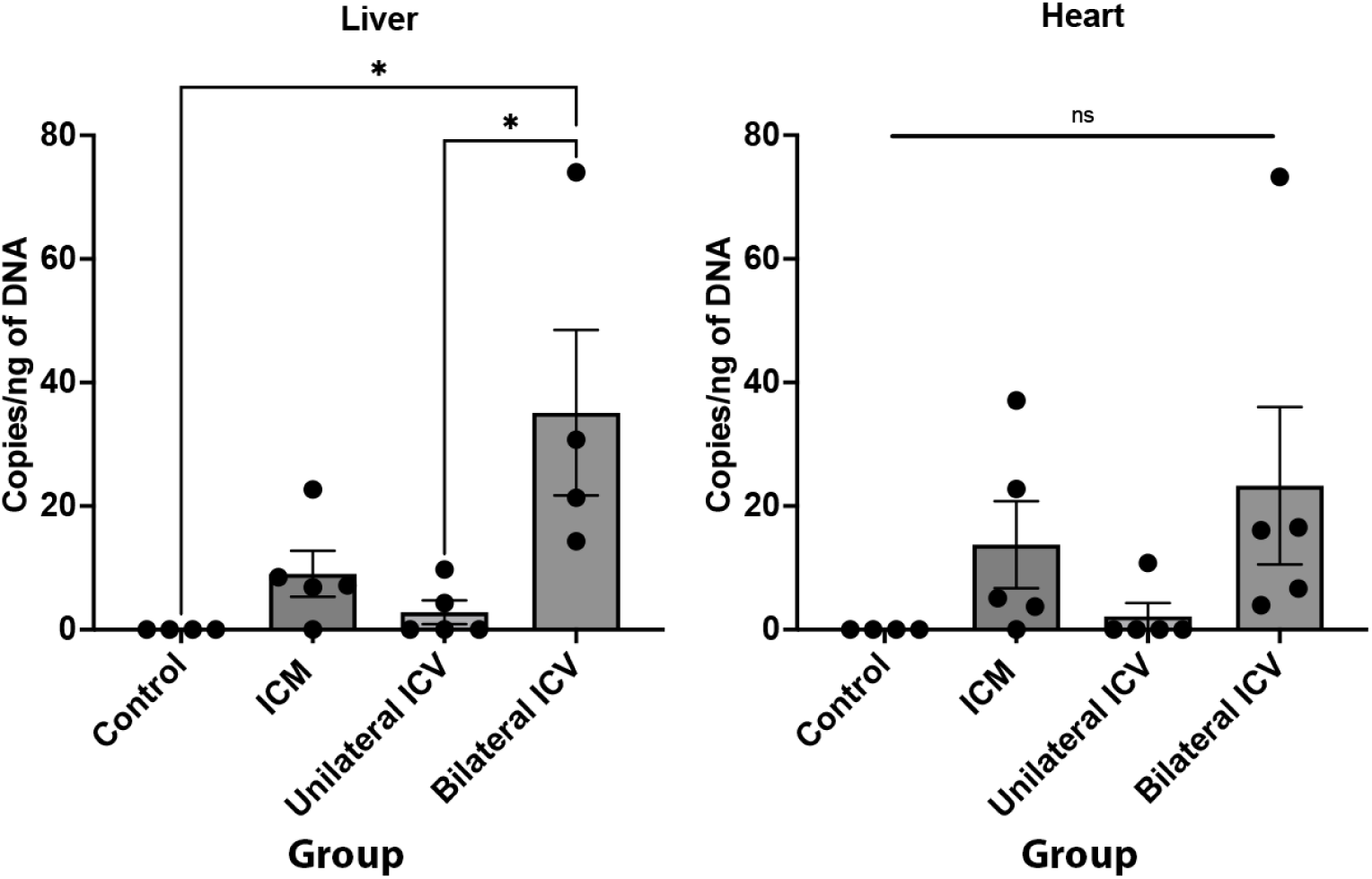
Quantification of viral copy number in liver and heart. qPCR on PNS tissue suggests that there is limited off-target transduction in peripheral organs. Note that Unilateral ICV achieves the lowest and most consistent number of viral copies in liver and heart. Also note that Bilateral ICV delivery results in significantly more viral copies in the liver. One-way ANOVA with Tukey’s correction for multiple comparisons. ns=not significant, *p≤0.05. Error bars = mean ± SEM.

**Table 8:** Viral copy number analysis in peripheral organs as determined by qPCR.

| Tissue | Number of viral copies/ng of DNA (Mean ± SEM) | | | |
| --- | --- | --- | --- | --- |
| | Control (n=4) | ICM (n=5) | Unilateral ICV (n=5) | Bilateral ICV (n≥4) |
| Liver | 0 | 9.0 ± 4 | 2.8 ± 2 | 35.1 ± 13 |
| Heart | 0 | 13.7 ± 7 | 2.2 ± 2 | 23.3 ± 13 |

When considering the best route of administration for motor neuron targeting, it is worth noting that a bilateral approach could lead to 10x more viral load in the liver than another delivery method, despite achieving similar levels of motor neuron transduction in spinal cord and brainstem. Overall, achieving a balance between specific transduction in the CNS versus off-target effects is a critical consideration.

## 4 Discussion

In line with previous studies, we have demonstrated that CNS-directed delivery of scAAV9-CMV-eGFP in neonatal mice can selectively transduce motor neurons to high levels. To our knowledge, we have presented evidence of the highest percentage of motor neuron targeting with a single injection of AAV9 to CSF in mice that has been published thus far, with unilateral ICV delivery of scAAV9-CMV-eGFP transducing over 88% of motor neurons in the lumbar spinal cord and over 69% of motor neurons in brainstem. More generally, at 4 weeks post-injection, we found that any CNS-directed route transduced over 68% of motor neurons in the lumbar spinal cord and over 55% of motor neurons in the brainstem. Viral copy numbers in the CNS were also high across all groups, however we noted that bilateral ICV delivery, whereby the titre is effectively doubled across two days, resulted in significantly higher viral copies in the liver. We also noted that bilateral ICV delivery leads to significantly more astrocytic transduction in spinal cord (around 1.4%) and brainstem (around 0.37%) than other delivery methods, although numbers remained low. In cortex, either unilateral or bilateral ICV approaches transduced significantly more astrocytes (>16%), compared to the ICM group, where astrocytic transduction was almost negligible.

Taking motor neuron transduction levels and viral copy number data together, unilateral ICV delivery may provide the best balance between achieving high and consistent lower motor neuron transduction whilst minimising peripheral effects. If astrocytic transduction is desired, then ICV approaches could be effective, but titre should be optimised to determine potential dosing thresholds that lead to astrocytic transduction with minimal peripheral impact. If avoiding astrocytic targeting, then an ICM approach may be more appropriate. Systemic delivery should also be considered if non-neuronal subtypes are the intended target, as has been shown previously in the literature. Further work is required to optimise targeting of upper motor neurons with these methodologies.

### 4.1 Systemic versus CNS-directed delivery of AAV9

Vascular delivery of AAV9 remains the most widely adopted infusion route and is also more applicable to paediatric patients. Whilst an intravascular approach has proven clinical potential and application, challenges remain. The most obvious challenge lies in the large quantities of vector that are required to achieve significant CNS-targeting since only a fraction of the vector penetrates the blood brain barrier (BBB), thus making widespread CNS delivery historically challenging (Foust et al., 2009, Gray et al., 2011), although development of BBB-penetrant capsids has been ongoing and several recent BBB-penetrating AAV9 capsid variants have been engineered that show widespread CNS transduction (Deverman et al., 2016, Chan et al., 2017, Nonnenmacher et al., 2021). With an intravenous approach using scAAV9-CBh-GFP in neonatal mice, Foust et al., 2009 reported one of the highest levels seen with a systemic approach, reporting up to 61% motor neuron transduction in mice. However, vascular delivery also leads to widespread targeting of astrocytes throughout the spinal cord and brain, and, most notably, in peripheral organs (Foust et al., 2009, Gray et al., 2011, Foust and Kaspar, 2009). Similar peripheral organ targeting, and astrocytic transduction in mice are reported with AAV9 after systemic delivery throughout the literature (Schuster et al., 2014, Mattar et al., 2015, Miyake et al., 2011, Gray et al., 2011, Duque et al., 2009, Rahim et al., 2011). Whilst targeting of astrocytes may not necessarily be undesirable, as there is potential for astrocytes to deliver secreted transgene products to the CNS, the additional challenges of toxicity, immunogenicity and circulating antibodies with such large infusion volumes are problematic.

With systemic delivery approaches, large vector doses are required to reach the CNS due to full-body distribution. Increasing the viral load consequently increases peripheral organ exposure. Indeed, significant concerns over liver toxicity have emerged in both clinical and non-clinical settings in spinal muscular atrophy, Duchenne muscular dystrophy and myotubular myopathy (Hudry et al., 2023, Nat.Biotech, 2020, Chand et al., 2021, Hamilton and Wright, 2021, Hinderer et al., 2018, Whiteley, 2023).The liver is highly vascularised, receiving around 25% of cardiac output and around 30% of the body’s total volume per minute (Abdel-Misih and Bloomston, 2010, Racanelli and Rehermann, 2006), which may drive hepatic tropism. The mechanisms by which the liver tolerates, or fails to tolerate, AAV9 have been reviewed recently (Keeler et al., 2019), however in brief transduction of AAV vector in macrophages in the liver, known as Kupffer cells, are thought to elicit an immune response and subsequently eliminate the therapeutic effect of delivered therapies by developing antibodies to the capsid, transgene, or through cytotoxic T-cell-mediated mechanisms. Activation of capsid-specific T cells may result in destruction of transduced hepatocytes and result in consequent hepatotoxicity. Acute liver injury has been found to correlate with high hepatocellular load, macrophage activation, and type 1 interferon innate virus-sensing pathway responses (Hudry et al., 2023). Thus, reducing the viral load, as can be achieved with CNS-directed AAV9 delivery, has clear advantages, despite being more invasive and technically challenging. The presence of anti-AAV antibodies in the bloodstream, even at low titres, can block transduction of tissues by intravenous AAV9 (Gray et al., 2011). Anti-AAV antibodies are common in human and non-human primates, although normally lower in infants and young children (Boutin et al., 2010, Calcedo et al., 2011, Samaranch et al., 2012).

Since reduced doses of vector are required to reach clinically relevant transgene expression with CNS-directed delivery, the immunogenicity risk is inherently lower. Thus, CNS-directed approaches offer a compelling alterative. Our data indicate that unilateral ICV delivery results in minimal viral genome copies in the liver, which is promising from a toxicity perspective. Notably, bilateral ICV delivery with an increased dose resulted in significantly higher viral load in the liver, however toxicity was not expected or observed with this viral load. Our observations may be due to increased titre of the vector, since we doubled the dose with a bilateral approach, however we cannot rule out other mechanisms of augmentation, which we will discuss. ICV delivery is already used in clinical trials such as those in Canavan disease (NCT04833907) and mucopolysaccharidosis type II (NCT04571970), and so is already clinically translatable. Previous work by has demonstrated around 41% motor neuron targeting in spinal cord with an ICV approach, however this work used a single stranded vector, which take longer to express than self-complementary vectors (Duque et al., 2009). Due to the close anatomical relationship between brainstem and the cisternal space, ICM routes of delivery in mice carry higher procedural risk. In the clinic, an intrathecal catheter via the subarachnoid space from a lumbar puncture may reduce invasiveness of an ICM procedure; in non-human primates, delivery to the lateral ventricle or ICM with a catheter has been shown to effectively target cervical motor neurons, although lumbar motor neuron transduction was not assessed (Kumagai et al., 2023). Our data suggest the main considering when choosing between an ICM or ICV route in murine preclinical work likely resides in desire to achieve astrocytic transduction. ICV appears to result in higher and more consistent lumbar motor neuron transduction, however, higher numbers of astrocytes are transduced, especially in cortex. If seeking to avoid astrocytic transduction, ICM may be a better option.

It is important to note that delivery of AAV9 into CSF directly does not completely negate the challenge of circulating pre-existing anti-AAV antibodies (Samaranch et al., 2012), and mild dorsal root ganglion toxicity has been highlighted in preclinical animal studies (Hinderer et al., 2018), however only one case has been reported in humans (Mueller et al., 2020). Since transduction can be achieved at lower viral titre within the CNS compartment, immunogenicity risks are reduced. Immune tolerance of AAV-based therapies, screening for neutralising antibodies and immunosuppression remain important clinical considerations. In addition, intra-CSF delivery of AAVs has been shown to induce more dorsal root ganglion (DRG) pathology in non-human primates than if AAV is delivered intravenously (Hordeaux et al., 2020). We did not assess DRG biodistribution here, however including this in future may allow us to better understand and predict the consequences of DRG transduction.

Note that we trialled intramuscular delivery of vector into gastrocnemius muscle in neonatal and adult mice, however, despite evidence of myofiber transduction, there was minimal evidence of retrograde transduction to motor neurons in the lumbar spinal cord (see appendix Fig A, 3). Previous studies have similarly shown that there is minimal transduction to lumbar spinal cord after direct intramuscular hindlimb delivery of AAV9 (Jan et al., 2019, Foust et al., 2009). This route of administration could be useful if seeking to selectively target peripheral muscle but does not appear useful for targeting alpha-motor neuron somata. It would be interesting to assess alternative approaches for evaluating CNS biodistribution. Co-injection of the same AAV serotype by two different routes with different reporters for each route of administration, for example, may provide an interesting way to compare distribution within the same animal. However, consideration of tolerable volumes, system pressure and distribution to other cell types, and visualisation of co-stains, would likely be pertinent. Assessing if both CSF-delivery methods remain equally as effective if vectors are delivered to mice during adulthood would be interesting, since age-dependent declines in transduction have been described with CNS-directed therapies (Garza et al., 2024). We chose to administer at neonatal stages as this is the major timepoint used in studies for therapeutic proof of concept and particularly where neurodegenerative phenotypes develop rapidly (Iannitti et al., 2018).

Another critical consideration is that here, we failed to find evidence of motor neuron transduction in the motor cortex. There could be several reasons for this linked into dose, route, age at time of delivery, and viral vector identity that merit further work. This is undoubtedly a key consideration for gene therapy work in diseases such as ALS, since both upper and lower motor neurons are affected. Our work highlights the importance of assessing and optimising viral distribution for preclinical work, to ensure vectors are reaching their intended target to ensure that they are assessed for true therapeutic efficacy.

### 4.2 Augmentation of transduction with consecutive delivery of AAV9

To achieve our highest titre group, we delivered vector across two sessions on consecutive days. The reasoning for this was primarily driven by our aim to increase vector titre within the constraints of the injection volume, to limit the risk of increasing intracranial pressure, whilst keeping the construct constant. At 4 weeks post-injection, we anticipated strong expression of the transgene (Hollidge et al., 2022). Fortuitously, this dosing regimen led to an interesting observation: when scAAV9-CMV-eGFP was delivered bilaterally across two days, effectively doubling viral titre, we noted that there was a significant increase in viral copy number in spinal cord and brainstem above the two-fold level that we might have anticipated. The mechanisms underpinning such an increase in viral vector load remain unclear.

The CNS is described as an immune-privileged compartment, which likely serves to preserve tissues with limited regenerative capacity from destructive inflammatory reactions (Forrester et al., 2018). In studies delivering small amounts of AAV vectors directly to the brain in Parkinson’s disease (Kaplitt et al., 2007, Marks et al., 2008, Marks et al., 2010, Christine et al., 2009), Canavan disease (McPhee et al., 2006, Leone et al., 2012) and late infantile neuronal ceroid lipofuscinosis (Worgall et al., 2008), AAV vector administration was generally associated with minimal detectable immune responses to the capsid or the transgene in serum and peripheral blood mononuclear cells. The upper limit of vector dose that can be administered without breaking the immunologic unresponsiveness of the CNS is unclear, but we might hypothesise that our initial dose increased host cell permissiveness, without initiating a substantial immune response.

The mechanisms of AAV transduction have been reviewed previously, with N-linked galactose described as the primary receptor for AAV9 along with binding of a putative integrin and laminin receptors as co-receptors (Shen et al., 2011, Bell et al., 2012, Dhungel et al., 2021). In brief, when AAV9 reaches a cell, it attaches to receptors/co-receptors and is endocytosed into the cell. Inside the endocytic vesicle, the virion travels towards the trans-Golgi network. During this trafficking phase, it goes through structural changes that expose a catalytic domain on the capsid. The virion then escapes into the cytosol and the capsid is imported through the nuclear pore complex into the nucleus in an importin B-dependent process. Here, the single stranded genome is ultimately released from the capsid. After release, it is converted to double stranded DNA and it persists as a circular episome, and linear or episomal concatemers. It is this final step of AAV9 transduction that permits expression of the transgene. Conversion into double stranded DNA (dsDNA) is described as the rate-limiting step for AAV9 transduction, that restricts the efficiency, speed of onset and transgene expression, since dsDNA conversion is independent of host-cell DNA synthesis and vector concentration (Hauck et al., 2004, Fisher et al., 1996, Ferrari et al., 1996). Augmentation of viral transduction can be achieved by improving the efficiency of double stranded DNA conversion, independently of vector titre. Current methods for this include using DNA-damaging agents such as ultraviolet irradiation or hydroxyurea, or inhibition of specific host-cell factors. However, the clinical utility of such approaches is limited, due to the risk of mutagenic breaks and exacerbation of genotoxicity. Further work is required to elucidate if our dosing approach may have influenced the cellular machinery and/or mechanisms of double stranded conversion, or influenced proteasomal degradation pathways, to consequently enhance transduction. Indeed, we might reasonably hypothesise that a threshold exists whereby higher doses reach a level that is sufficient to transduce glial cells, however our finding of significant cortical astrocyte transduction following ICV administration may indicate other mechanisms at play.

Overall, further work is required to understand the complex mechanisms by which repeated AAV9 delivery across consecutive days can increase viral load in the spinal cord, and the utility, if any, of such an approach.

## Acknowledgments

We acknowledge the expertise of technical staff within the SITraN histology lab of University of Sheffield for their assistance. We also acknowledge the help of animal technicians within facilities at the University of Sheffield for their care and assistance with monitoring pregnant dams. This work was supported by the MND Association and MND Scotland [2023/MNDS/6501/791MEA].

## 5 Author Contributions

A.J. Mole: Data curation, formal analysis, investigation, methodology, project administration, software, supervision, visualisation, writing – original draft, writing – review & editing

C.F. Sander: Data curation, formal analysis, methodology, software, visualisation, writing – review & editing

A.R. Parmar: Methodology, validation, visualisation, writing – review & editing

A.J. Wlliams: Methodology, visualisation, writing – review & editing

1. M. Azzouz: Conceptualization, funding acquisition, supervision, writing – review & editing

G.M. Hautbergue: Conceptualization, funding acquisition, supervision, writing – review & editing

P.J. Shaw: Conceptualization, funding acquisition, supervision, writing – review & editing

1. L. Ferraiuolo: Conceptualization, funding acquisition, supervision, writing – review & editing

R.J. Mead: Conceptualization, funding acquisition, project administration, resources, supervision, writing – review & editing

## List of Abbreviations

Abbreviation: Expansion
AAV: Adeno-associated virus
ALS: Amyotrophic lateral sclerosis
BBB: Blood brain barrier
CHAT: Choline acetyltransferase
CMV: Cytomegalovirus
CSF: Cerebrospinal fluid
GFAP: Glial fibrillary acidic protein
GFP: Green fluorescent protein
IBA1: Ionised calcium binding adaptor molecule 1
ICM: Intra-cisterna magna
ICV: Intra-cerebroventricular
IHC: Immunohistochemistry
IT: Intrathecal
IV: Intravenous
MN: Motor neuron
SMA: Spinal muscular atrophy
VG: Viral genomes

## 9 Appendices

**Fig. A. 1:**
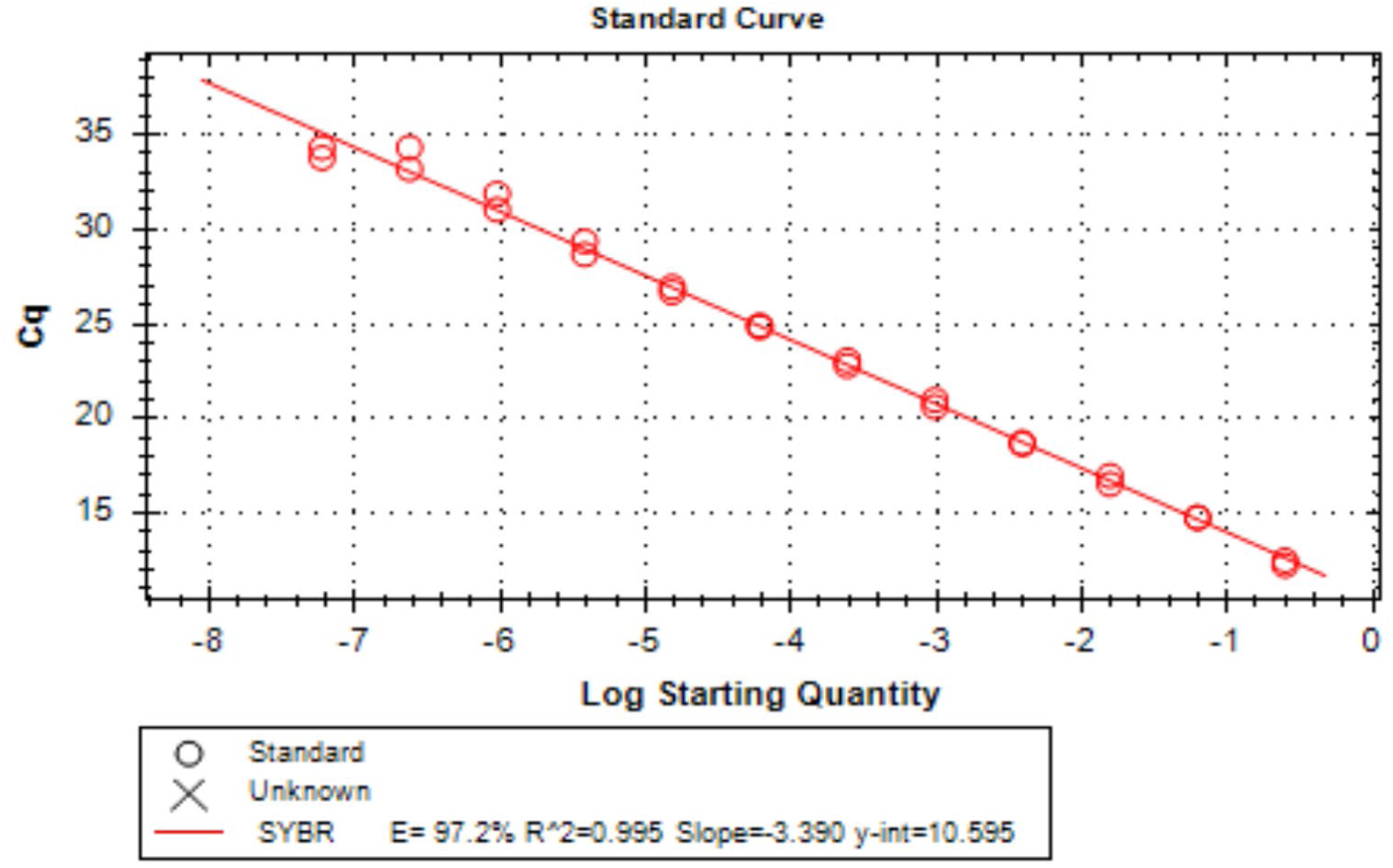
**Standard curve of CMV.GFP plasmid clone used for quantitative determination of viral genome copy number in central tissues by qPCR.**

**Fig. A. 2:**
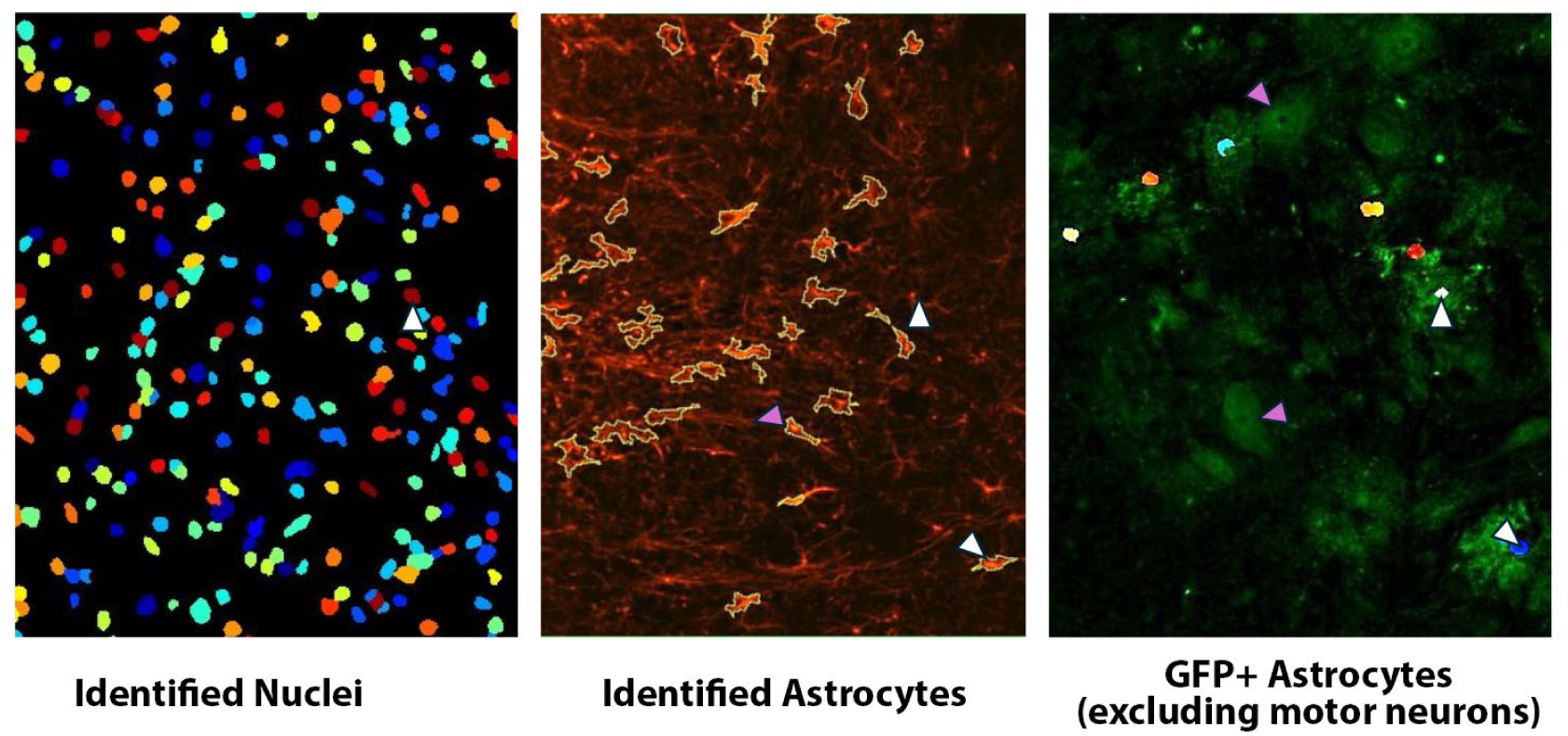
Cell profiler mask examples showing identification of astrocytes with application of machine learning tool. Note example showing the identification of an astrocyte with nucleus that co-expresses GFP (white arrow). Also note that although the motor neurons (pink arrows) express GFP, the machine was trained to exclude these from quantification.

**Fig. A. 3:**
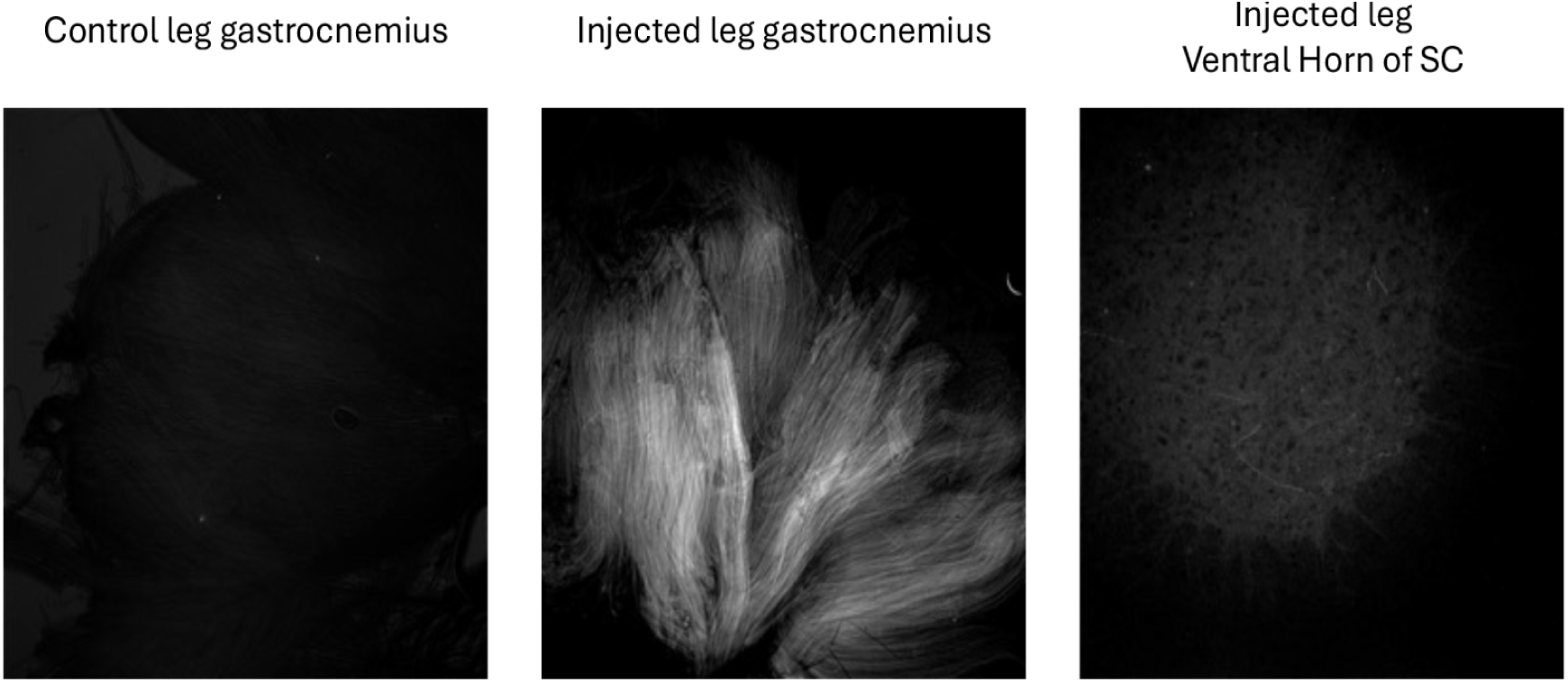
Immunohistochemistry after intramuscular delivery of AAV9-CMV-eGFP suggested that, despite transducing myofibers, there was minimal retrograde transduction of AAV9, as GFP was not detected in lumbar spinal cord motor neurons. Previous studies have similarly shown that there is minimal transduction to lumbar spinal cord after direct intramuscular hindlimb delivery of AAV9 (Jan et al., 2019, Foust et al., 2009). Intramuscular delivery of AAV9 could be useful if seeking to selectively target peripheral muscle.

